# Design of a sequestration-based network with tunable pulsing dynamics

**DOI:** 10.1101/2024.03.24.586474

**Authors:** Eiji Nakamura, Christian Cuba Samaniego, Franco Blanchini, Giulia Giordano, Elisa Franco

## Abstract

Incoherent feedforward networks exhibit the ability to generate temporal pulse behavior. However, exerting control over specific dynamic properties, such as amplitude and rise time, poses a challenge and is intricately tied to the network’s implementation. In this study, we focus on analyzing sequestration-based networks capable of exhibiting pulse behavior. By employing time-scale separation in the fast sequestration regime, we approximate the temporal dynamics of these networks. This approach allows us to establish a mapping that elucidates the impact of varying the kinetic rates and pulse specifications, including amplitude and rise time. Furthermore, we introduce a positive feedback mechanism to regulate the amplitude of the pulsing response.

## I. Introduction

Biological systems use the temporal dynamics of various parameters, such as concentration, enzymatic activity, allosteric configuration, biochemical modification, or spatial localization, to interpret and respond to external stimuli [1], [2]. In this context, transient pulses are a prevalent type of temporal pattern found across biology. Cells can encode information into several temporal features of pulses, such as amplitude, duration, and frequency of pulses, and this encoding can enable a variety of functions [3]–[6]. From an engineering perspective, the adoption of pulsed signals has the potential to expand the capacity of synthetic biological circuits to store and transmit information using a limited number of molecular components [7]–[10].

Pulse generation in biomolecular systems can be realized by specific circuit motifs, such as the incoherent feedforward loop (IFFL) [11], [12], and negative feedback [13], which present nonlinear dynamics. However, from a mathematical standpoint, the easiest way to generate a pulse is through the subtraction of two exponential functions [14] (Fig. 1). This suggests that a chemical reaction network capable of implementing a subtraction operation could be used to build a pulse generator. Chemical reaction networks that perform subtraction have been demonstrated through molecular sequestration, a versatile motif that is also central for the construction of biomolecular integral controllers [15]–[18]. Sequestration occurs when two chemical species *A* and *B* interact via a second order reaction 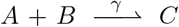. If the sequestration reaction rate parameter *γ* is sufficiently large, and if timescale separation arguments can be applied, then this reaction can be used to compute the difference between the concentration of *A* and of *B* [17]. Here we take advantage of fast sequestration and its capacity to subtract signals to build a pulse generator circuit, whose topology is comparable to the topology of an IFFL [19]. We rigorously show that the dynamics of this pulse generator can be approximated well as the difference of two exponential functions. The approximated solution also enables us to analytically derive some features of the pulse, such as amplitude and peak timing. Finally, we show that the amplitude of pulse signals can be enhanced by including positive feedback to our sequestration-based pulse generator.

**Fig. 1.**
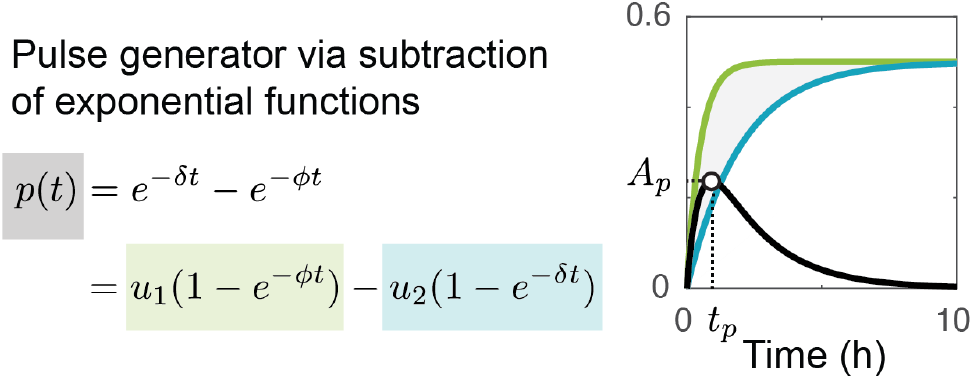
A pulse can be generated by the subtraction of two exponential functions.

## II. Subtraction for pulse generation

We can generate a pulse by the subtraction of two functions: (*e*^−*ϕt*^ − *e*^−*δt*^), with distinct exponential parameters *ϕ* and *δ* [14]. To create an intuition of how this subtraction produces a pulse, we rewrite it as (1 − *e*^−*δt*^) − (1 − *e*^−*ϕt*^). Further, each function can be scaled by gains *u*_1_ and *u*_2_, yielding the difference *p*(*t*) = *u*_1_(1 − *e*^−*δt*^) − *u*_2_(1 − *e*^−*ϕt*^). As an example, in Fig. 1 we show that if *u*_1_ = *u*_2_ = 0.5, and *ϕ* < *δ*, the first function (green) converges to steady state faster when compared to the second function (blue). For this reason, their difference *p*(*t*) (gray region) is non-negative at all times, and it first increases and then decreases. As a result, function *p* exhibits a pulsed behavior (black line).

The expression of *p*(*t*) allows us to find the peak time, *t*_*p*_, and the peak amplitude, *A*_*p*_ = *p*(*t*_*p*_), of the pulse, by solving 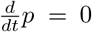. This results in 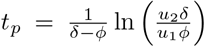, and 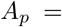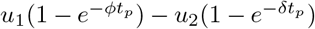. Therefore, specifications on the pulse peak time and amplitude can be met by a molecular network whose behavior approximates the subtraction of two exponential functions. These specifications depend directly on the time constants.

## III. A Sequestration-Based Pulse Generator

We use molecular sequestration to implement the subtraction of two biochemical signals to generate a pulsed output behavior as shown in Fig. 2. Our chemical reaction network includes two species *X* and *Z* produced by a zero order reaction; for simplicity, we assume the same production rate parameter *u*. Species *Z* then catalytically produces species *Y* with rate constant *β*. Species *X* and *Y* then sequester each other, through a second order reaction with rate constant *γ*, to produce *C*. Finally, species *X, Y* and *C* decay at rate *δ*, while species *Z* decays at rate *ϕ*. We list below a summary of all the chemical reactions:

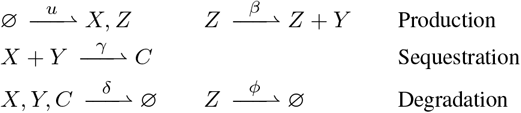

**Fig. 2.**
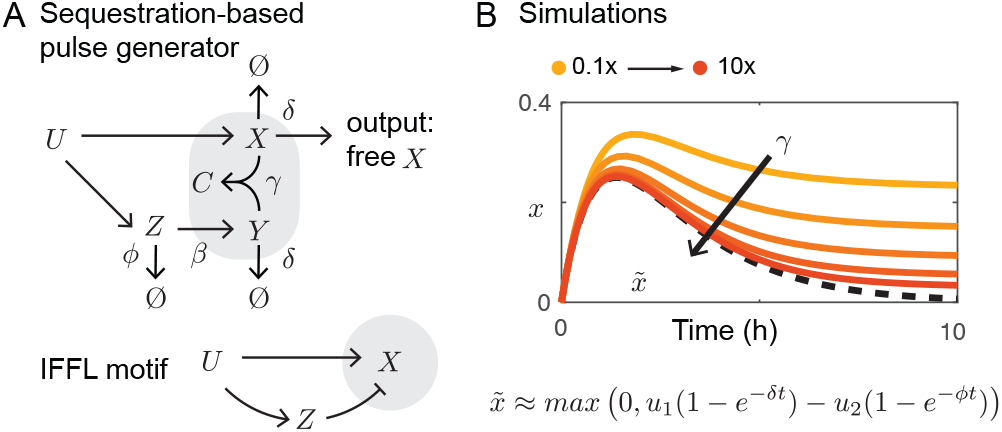
A: Chemical reaction network implementing a pulse generator, compared to IFFL topology. B: Simulations comparing the approximated value 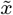 (dashed black line) with the non-approximated output *x* for different values of the sequestration rate constant *γ*.

We use the law of mass action to derive a set of Ordinary Differential Equations (ODEs) that describe the dynamical time evolution of the species concentrations:

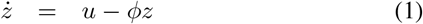

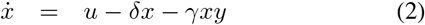

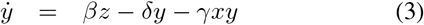

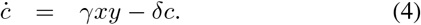

Because species U directly generates *X*, as well as indirectly removes *X* through *Y*, the structure of this chemical reaction network is akin to the topology of an IFFL, as illustrated in Fig. 2 A.

## IV. Pulse Generator Dynamics: Stability AND Exponential Approximation

We start by proving the following stability property.

### Proposition 1.

*For u* > 0, *system* (1)-(4) *admits a unique positive equilibrium, which is globally asymptotically stable*.

*Proof*. Equation (1) implies that z(*t*) converges exponentially to its equilibrium value 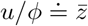. Consider now (2)-(3) with constant input 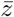, the steady state from (1). To show that an equilibrium exists, set 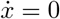 and 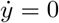

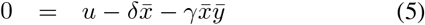

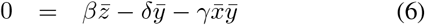

to derive 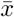 and 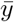. By taking the difference 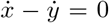, we get

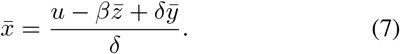

Then, replacing 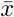 in (6) yields the second order equation

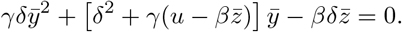

Take 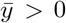 as the only positive root of this equation and then find 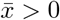 from equation (7). To prove global stability, once 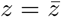, let us subtract (5)-(6) from (2)-(3) to write the x-y system as

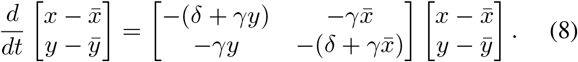

Since the matrix appearing in (8) is column diagonally dominant, it admits the 1-norm as Lyapunov function (as it can also be computed through the algorithm in [20], [21]), which proves global stability of the x-y subsystem (2)-(3). Finally, from (4), since 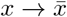 and 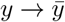, then 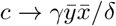, the only possible equilibrium value.

To analyze the transient dynamics in the regime of fast sequestration (*γ* → ∞), we adopt the change of variables *w*_1_ = *x* + *c* and *w*_2_ = *y* + *c*, and rewrite the model as

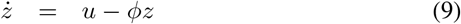

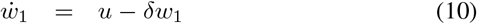

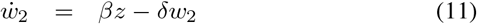

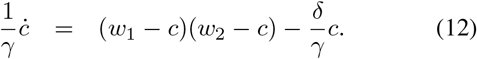

Following a similar analysis as in previous work [17], [18], we obtain the approximations *c* ≈ min(*w*_1_, *w*_2_) and 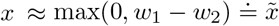. These approximations are evident in computational simulations, as shown in Fig. 2B (simulation parameters are listed in Table I), and can be formally proven as follows.

### Proposition 2.

*Assume that u* > 0 *is a step input and all variables are initialized at 0: w*_1_(0) = *w*_2_(0) = *z*(0) = *c*(0). *Then, as γ* → ∞, *the corresponding solution c*_*γ*_ *of* (12) *converges to* min(*w*_1_(*t*), *w*_2_(*t*)) *from below, uniformly with respect to γ*.

*Proof*. We perform an infinitesimal analysis at time zero, looking for the first nonzero derivative of each variable. At *t* = 0, 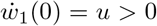, while 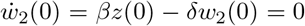 And 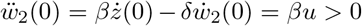. Moreover, we have

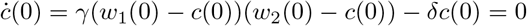

and

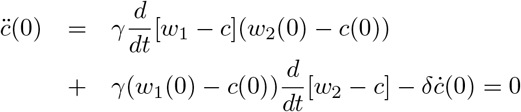

Since the first non-zero derivative of *w* _1_ is its first derivative 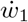 and the first non-zero derivative of *w* _2_ is its second derivative 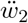, and since the first and the second derivatives of *c* are 0, in a right neighborhood of 0, *t* ∈ [0, ϵ], it must be *c*(*t*) ≤ *w*_1_(*t*) and *c*(*t*) ≤ *w*_2_(*t*).

The next step is to show that these inequalities are valid for all *t*. Assume by contradiction that *c* surpasses the minimum of *w*_1_ and *w*_2_ from below: at some 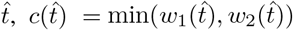. Then either 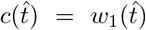 or 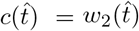. Consider the derivative,

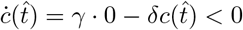

On the other hand, both 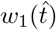 and 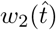 have nonnegative derivatives: hence, in a right neighborhood of 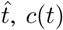 would be smaller than both 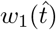 and 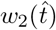. This is a contradiction. Therefore, *c*(*t*) can surpass neither *w*_1_ nor *w*_2_.

The final step is to show that we have uniform convergence of *c*_*γ*_ (*t*) to the function 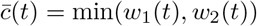. We have to notice that 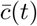 is non differentiable in general, hence we need to consider its right Dini derivative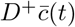, which is uniformly bounded as

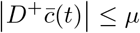

for some *μ* > 0, because *w*_1_ and *w*_2_ behave as the step response of a stable linear system.

We prove that for any small *ϵ* > 0, there exists a 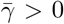 such that, for 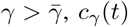 is lower and upper bounded as

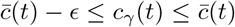

We have already proven that 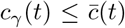. Hence, we need to prove that 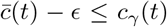 or, equivalently, defining the function 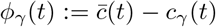, that

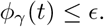

By continuity, the condition is true in a right neighbourhood of 0, 0 ≤ *t* ≤ *δ*, because both 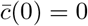 and *c*_*γ*_ (0) = 0, hence *ϕ*_*γ*_ (0) = 0. Now we show that, for *γ* > 0 large enough, if 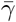 is large, then *ϕ*_*γ*_ (*t*) remains below *ϵ* for all *t* > 0 (and not just for 0 ≤ *t* ≤ *δ*, as we have seen so far). Assume by contradiction that this is not the case, namely, that function *ϕ*_*γ*_ (*t*) grows over *ϵ*: for some 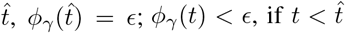; and *ϕ*_*γ*_ (*t*) > ϵ, in a right neighborhood of 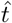. We show that this is not possible because the derivative of *ϕ*_*γ*_ (*t*) is negative at 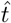, for large *γ*. The condition 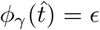 implies that 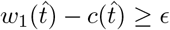 and 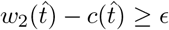, because 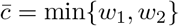. Consider the derivative of 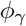:

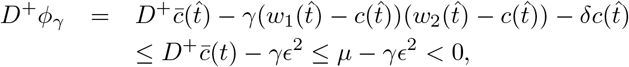

provided that 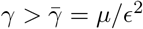. Since the derivative is negative, *ϕ*_*γ*_ cannot grow above *ϵ*.

Then, for arbitrary small *ϵ*, we can take 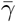 large enough so that *ϕ*_*γ*_ ≤ *ϵ* for 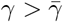.

From the equations for 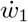 and 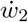, we can derive the expressions in terms of Laplace transforms:

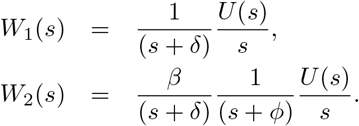

By anti-transforming, in the time domain we get

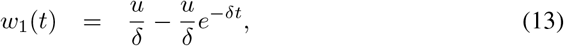

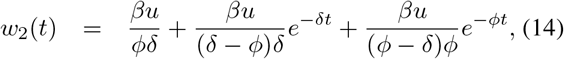

and their difference can be written as

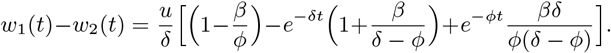

Hence, 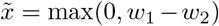 admits the explicit expression

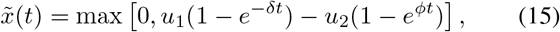

where 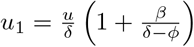 and 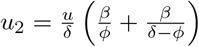.

### Remark 1.

*Proposition 2 implies that x*(*t*) *converges to* max(0, *w*_1_ − *w*_2_) *as γ* → ∞.

The main mechanism driving the circuit behaviour, illustrated in Fig. 3, is the following. From equations (13) and (14), as discussed in the proof of Proposition 2, we see that *w*_1_(0) = *w*_2_(0) = 0 and, for small times *t* > 0, *w*_1_(*t*) > *w*_2_(*t*), since 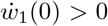 while 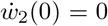 (this can be also verified via the initial limit theorem of the Laplace transform). If the asymptotic value for *w*_2_ is larger than the asymptotic value for *w*_1_, i.e. if the system parameters satisfy the inequality

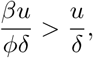

namely, *β* > *ϕ*, then *w*_2_(*t*) must become larger than *w*_1_(*t*) after some time 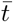. Hence, 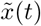 is positive for 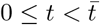 and is zero for 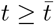.

**Fig. 3.**
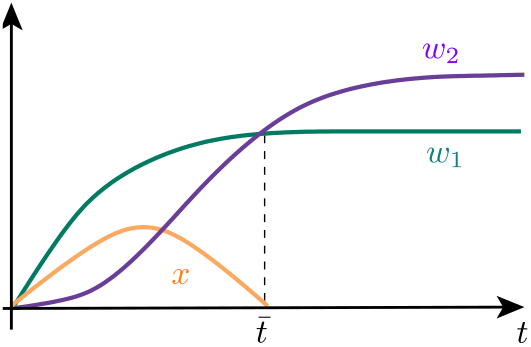
The approximation of the output 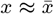 holds when (*w*_1_ − *w*_2_) is positive

## V. Pulse Dynamics Specifications and Parameter Sensitivity

With simulations, in Fig. 2B we compare the approximated solution 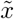 (dashed black line) and the full solution of *x* for different values of sequestration rates *γ* (parameters listed in Table I). Larger values of *γ* make the full solution x converge to the approximate solution 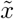.

To further test the capacity of the approximated solution 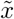 to capture the dynamics of the system, we examine the solution when the parameters are varied with respect to those in Table 1, as shown in Fig. 4A. We use equation (15) to compute 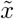 as a function of the parameters. Parameter *β* influences the relative magnitude of the approximation gains *u*_1_ and *u*_2_: when *β* > *ϕ*, this results in *u*_2_ > *u*_1_, and makes the pulse’s basal level converge to very low values. When *β* < *ϕ* (therefore *u*_1_ > *u*_2_), the basal level is larger than zero, compromising the pulse behavior. If *δ* < *ϕ*, we observe a well defined pulse behavior. When *δ* increases (approaching *ϕ*), the pulse behavior disappears. Picking *ϕ* > *δ* appears to be sufficient to achieve a pulse behavior, but changes in *ϕ* also influence the gains. The ratio *β*/*ϕ* determines which gain is larger (*u*_1_ or *u*_2_), therefore *ϕ* has the opposite effect of *β*: a large *ϕ* (*β* < *ϕ*) would yield *u*_2_ < *u*_1_ and a loss of the pulse behavior. Finally, the input *u* affects both *u*_1_ and *u*_2_ but it does not determine which one is larger. The magnitude of *u* primarily influences the amplitude of the pulse.

**TABLE 1.** Parameters for the molecular sequestration-based pulse generator model.

| Parameter | Unit | Value |
| --- | --- | --- |
| $u = \phi = \beta = \delta/2$ | $\text{h}^{-1}$ | 0.5 |
| $\gamma$ | $\mu\text{M}^{-1}\text{h}^{-1}$ | 50 |

**Fig. 4.**
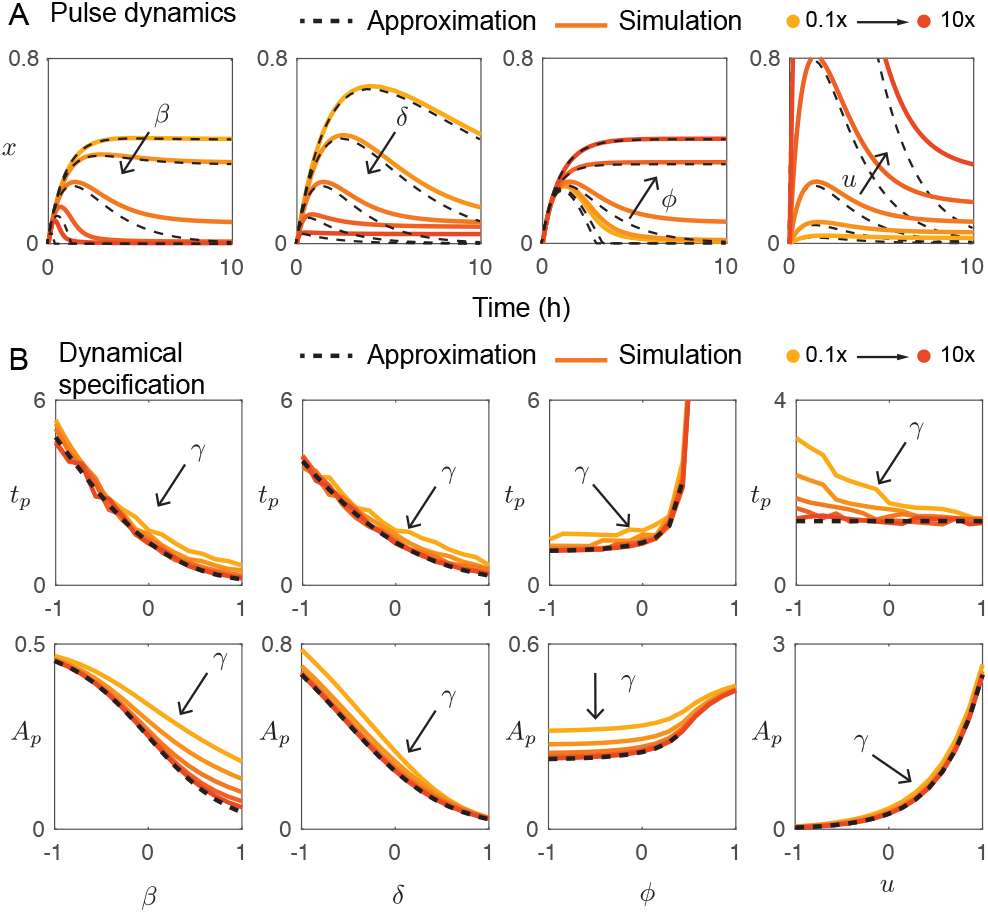
Molecular pulse generator. (A) Simulated sequestration (yellow-orange) is compared to the general approximation when generating a pulse (black dashed line). Production rates, degradation rates, and input concentrations are varied to test the effects on the approximations. (B) The amplitude *A*_*p*_ and the peak time *t*_*p*_ of the simulated (black dashed) and approximated (yellow-orange) pulses are compared for an increasing sequestration rate.

In Fig. 4 B we show that the approximated solution 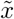 captures the effect of each parameter on the peak time *t*_*p*_ and the peak amplitude *A*_*p*_; the full solution is reported for comparison, as nominal parameters in Table 1 are varied. As long as *γ* is sufficiently large (fast sequestration) the approximate solution captures and predicts the peak time and amplitude.

## VI. A Positive Feedback Loop Controls THE Pulse Amplitude

Next, we investigate through simulations the effect of an additional positive feedback loop at the output node (*X*) of our pulse generator:

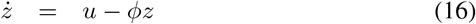

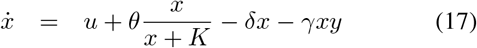

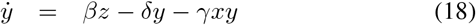

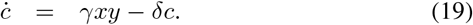

Term 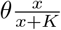 corresponds to a Michaelis-Menten approximation of an autocatalytic process, where *θ* and *K* represent the maximum production rate and the Michaelis constant of the positive feedback respectively. In Fig. 5 A, the new positive feedback loop is represented as a red arrow. We evaluate the effects of the positive feedback loop by comparing the original IFFL network (1)–(4) with the solution of the modified network (referred to as IFFL+PF). In these simulations we set parameter *ϕ* = 0.05; this choice makes the effects of the positive feedback loop more visible. Other parameters are left unchanged and are listed in Table 1.

**Fig. 5.**
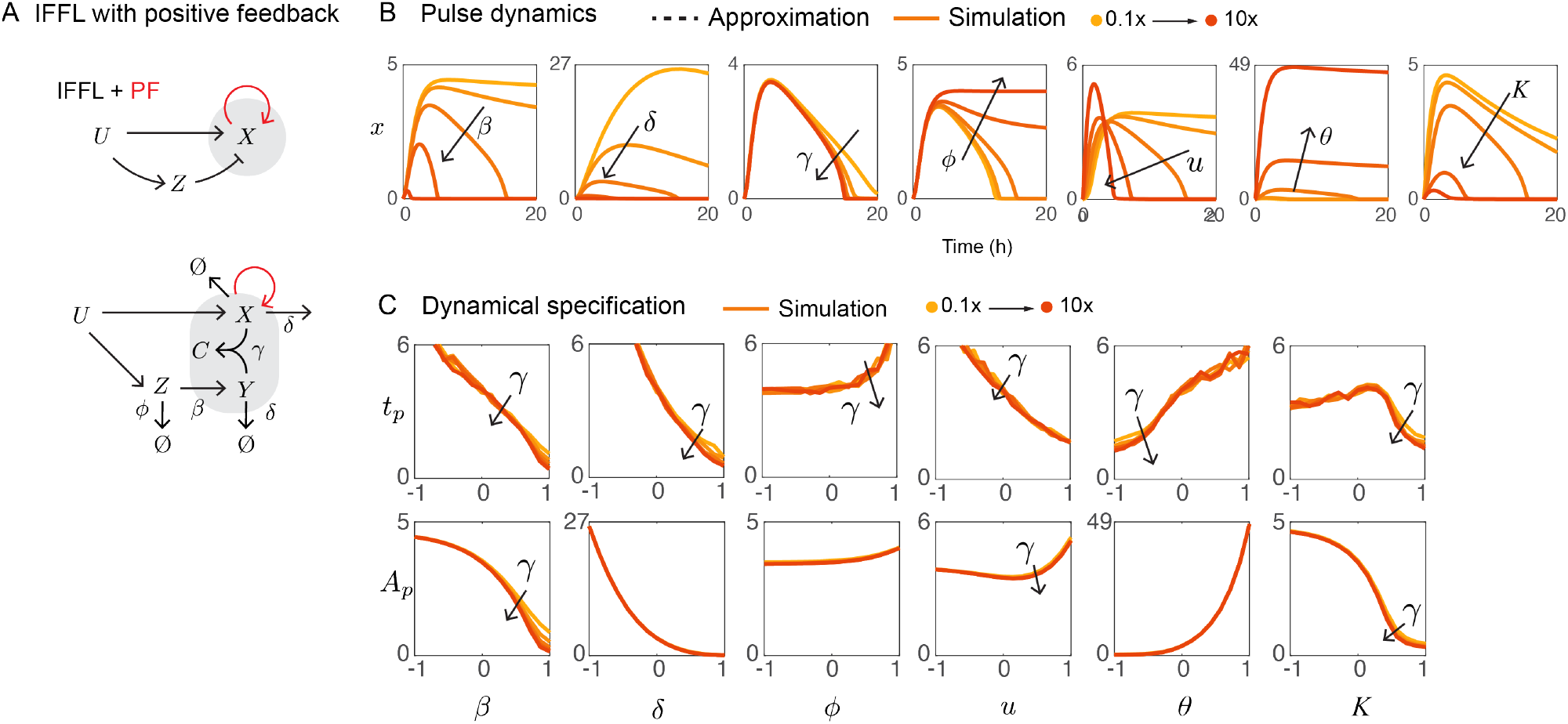
(A) The left diagram shows a general motif of incoherent feed forward loops (IFFL) with a positive feedback that we derived an similar motif utilizing molecular sequestration from in the bottom diagram. (B) Simulation results of IFFL with a positive feedback (yellow-orange) are shown. Production rates, sequestration rate, degradation rates, input strength, strengh and threshold of the feedback positive feedback are individually varied to test the effects of those parameters on the circuit dynamics. (D) The amplitude, *A*_*p*_ and the peak time *t*_*p*_ of the simulated pulses (yellow-orange) are shown with an increasing sequestration rate.

Fig. 5 B shows the circuit dynamics when individual parameters are varied. Compared with the original IFFL model, the amplitude and duration of IFFL+PF drastically changes, which suggests that the positive feedback provides the circuit with extended dynamic range of those properties. These can also be observed in Fig. 5B; when the ranges of the y-axes in Fig. 4B and Fig. 5 are compared, we note a wider amplitude range for the IFFL+PF dynamics, which is the most apparent under changes in parameters *δ* and θ. One noticeable difference from the original IFFL is that the curves for *t*_*p*_ and *A*_*p*_ are not always monotonic. With the original IFFL, *t*_*p*_ and *A*_*p*_ curves show the same trend against each parameter (except for *u*); for instance, both *t*_*p*_ and *A*_*p*_ increase as *ϕ* increases (Fig. 4B). This means that in the original IFFL it is hard to adjust *t*_*p*_ and *A*_*p*_ individually. On the other hand, for the IFFL+PF, the curves of *A*_*p*_ as a function of *u* and of *t*_*p*_ as a function of K do not monotonically increase or decrease: they have an extremum, which can be attributed to the additional non-linearity given by the positive feedback. The non-monotonic trends in the dynamic behavior of the IFFL+PF circuit make it more controllable and enable tuning *t*_*p*_ and *A*_*p*_ individually. It is worth noting that *t*_*p*_ and *A*_*p*_ did not show strong dependence on *γ*. That is because the elevated level of *X* leads to faster sequestration rate, which almost use up *Y* while *X* is present. Thus, the sequestration rate is limited by y rather than *γ*.

It should be noted that an increased amplitude can cause the loss of pulsatility. Here we define the pulsatility index of a signal as

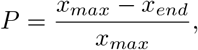

where *x*_*max*_ is the peak value of *X* (which is equivalent to the amplitude), and *x*_*end*_ is the value at the final time *t* = 20 [h]. Fig. 6B shows contour panels of the pulsatility index P (top) and amplitude *A*_*p*_ (bottom) on two-dimensional parameter planes. Although we can have parameter planes with any combinations of two parameters, here we are showing parameter planes with K versus each of the other parameters. In most of the parameter planes, comparing P and *A*_*p*_ panels of each parameter planes, the low-P area (for instance on the lower left area of the K-*β* plane) roughly coincides with the high-*A*_*p*_ area on the *A*_*p*_ panel. This means that the positive feedback loop not only increases the amplitude, but also increases the pulse duration, which ultimately results in the loss of pulsatility (P = 0). Therefore, in the parameter regimes evaluated here, there is a tradeoff between P and *A*_*p*_. While this trend is basically maintained in most of the parameter planes, K-*ϕ* and K-*u* planes have large overlap between the high-P area and high-*A*_*p*_ area, and the other planes also have small overlaps. Therefore, with the positive feedback, we can tune the amplitude of a pulse within a wider dynamic range.

**Fig. 6.**
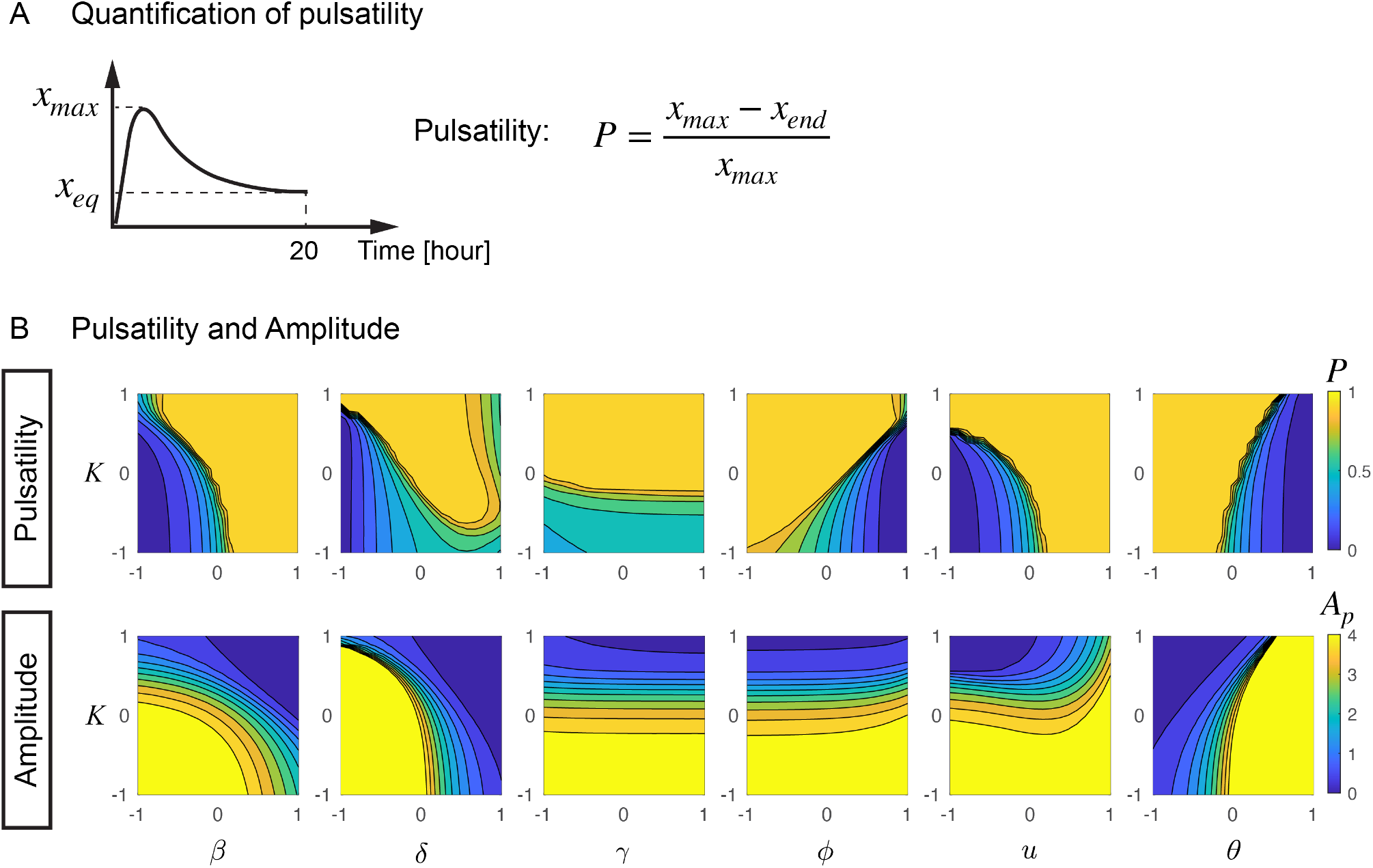
(A) A schematic of a pulse signal to visually explain the concept of pulsatility *P*. (B) Contour maps of pulsatility *P* (top) and amplitude *A*_*p*_ (bottom) on two-dimensional parameter planes are shown to evaluate the relationship between pulsatility and amplitude. All parameter planes have the same y-axis, *K*. The color scales are the same for all the contour maps; 0 to 1 for *P*, and 0 to 4 for *A*_*p*_ as shown on the right.

## VII. Conclusion

Pulse generation is one of the most prevalent phenomena in living systems, and also a basic function when designing circuits that are able to process temporal signals. We have discussed a subtraction-based pulse generator, and shown that the molecular sequestration mechanism can be used to build a molecular pulse generator in practice. Experiments *in vitro* demonstrated that a molecular realization similar to our system can generate tunable pulsatile behavior, as well as the ability to detect fold changes of the input [19]. We rigorously derived mathematical expressions to approximate the dynamics of the circuit in the regime of fast sequestration: the approximate solution of the pulse generator takes the form of a subtraction of two exponential functions. The approximate solution was validated through computational simulations that show convergence of the approximate solution to the actual solution as the sequestration rate increases. We finally suggested that positive feedback can be introduced to tune the behaviour of the pulse generator. Through simulations, we have shown that a positive feedback loop at the output node can enhance the amplitude of the signal, while maintaining a pulsatile behavior, so that the pulse generator has an increased dynamic range of the amplitude. However, the positive feedback loop may induce bistability, which can cause the loss of pulsatility. To quantitatively assess how each parameter affects pulsatility, we have proposed an index, P, and shown that, by carefully choosing the parameter values, the circuit endowed with the positive feedback can maintain a high P value with a larger dynamic range of the amplitude *A*_*p*_ with respect to the circuit without the positive feedback.

